# Developmentally stable representations of naturalistic image structure in macaque visual cortex

**DOI:** 10.1101/2024.02.24.581889

**Authors:** Gerick M. Lee, C. L. Rodríguez-Deliz, Brittany N. Bushnell, Najib J. Majaj, J. Anthony Movshon, Lynne Kiorpes

## Abstract

We studied visual development in macaque monkeys using texture stimuli, matched in local spectral content but varying in “naturalistic” structure. In adult monkeys, naturalistic textures preferentially drive neurons in areas V2 and V4, but not V1. We paired behavioral measurements of naturalness sensitivity with separately-obtained neuronal population recordings from neurons in areas V1, V2, V4, and inferotemporal cortex (IT). We made behavioral measurements from 16 weeks of age and physiological measurements as early as 20 weeks, and continued through 56 weeks. Behavioral sensitivity reached half of maximum at roughly 25 weeks of age. Neural sensitivities remained stable from the earliest ages tested. As in adults, neural sensitivity to naturalistic structure increased from V1 to V2 to V4. While sensitivities in V2 and IT were similar, the dimensionality of the IT representation was more similar to V4’s than to V2’s.

## Introduction

Behavioral performance improves during early life – animals learn new skills, and refine their performance within the space of skills they already know (Blumberg and Adolph, 2023; Dekker et al., 2020; Diamond and Goldman-Rakic, 1989; Kiorpes, 2016; Pistorio et al., 2006). The basis of these improvements is of interest – the acquisition of new skills often reflects a mixture of neural, muscular, and morphological changes (Adolph and Hoch, 2019), but improvement on psychophysical tasks can more simply be linked to changes in the brain. By understanding where these changes take place, we gain insight into how behavior emerges from the combined activity of sensory, association, and motor areas.

In the macaque visual system, behavioral capabilities on basic spatial vision tasks improve throughout the first year of life (Kiorpes, 2016). Despite this, neural sensitivity in the LGN, V1, and V2 is mature by roughly 16 weeks of age (Kiorpes and Movshon, 2004; Movshon et al., 2005; Zheng et al., 2007). Neurons in the inferotemporal cortex of infant macaques – like those in adults – can be selective for object identity, but are immature in their temporal dynamics (Rodman et al., 1993). Beyond this, the extent to which neural activity in developing V4 and IT limits behavioral development remains unknown. We wondered whether the gap between neural and behavioral development reflects immaturity in downstream visual areas like V4 or IT, or from more remote areas that convert sensory information into decisions and actions.

To address these questions, we used synthetic visual textures (Portilla and Simoncelli, 2000) (Figure 1A) whose structural similarity to natural images can be titrated (Freeman et al., 2013), and which preferentially drive activity in areas V2 and V4 (but not V1) (Freeman et al., 2013; Okazawa et al., 2015, 2017). We designed a 4 choice task which allowed us to quickly and reliably estimate the behavioral texture sensitivities of macaque monkeys across development. We then used multielectrode recording arrays to measure neural texture sensitivities at the single site and population level from areas V1, V2, V4, and the posterior portion of inferotemporal cortex (IT). We used the same images as in behavior, as well as a larger set of similar images (Figure 1B). These measurements allowed us to compare neural encoding with behavioral performance, in overlapping sets of animals, as a function of age. They also allowed us to relate how early, middle, and late ventral visual areas encode naturalistic textures at all ages, and to bridge our understanding of how sensitivities to naturalistic texture in areas like V2 and V4 might support information representation in area IT.

**Figure 1:**
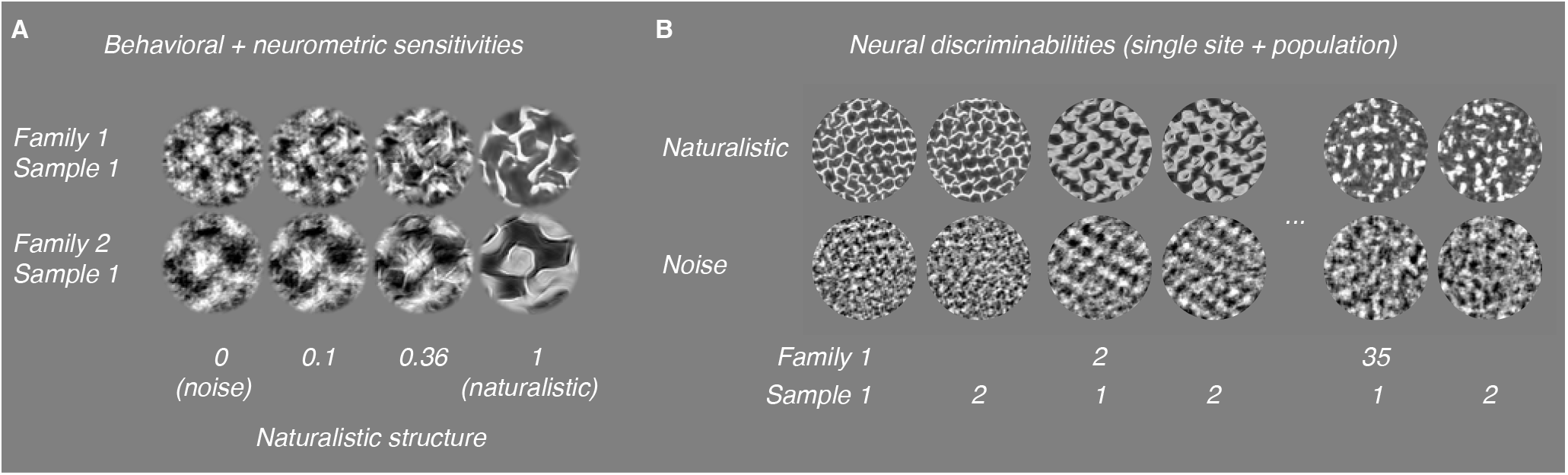
Naturalistic texture stimuli. A: Texture images varying continuously in the strength of their naturalistic structure. We used these images for measurements of behavioral and neurometric sensitivities. B: Naturalistic (top row) and noise textures (bottom row) used to characterize discriminability. Here, we used a total of 35 families, each containing 15 samples.

## Results

We made behavioral measurements of naturalistic texture sensitivity from 7 macaques, including 5 tested longitudinally from as early as 16 weeks, through the first 1-2 years of life, as well as 2 older controls. We made neural measurements of naturalistic texture encoding from 6 animals in total – 5 during the first year of life, as well as 2 adults (we implanted one animal twice).

### Behavioral sensitivities double during the first year of life

To measure animals’ behavioral sensitivities, we designed a 4 choice oddity task (Figure 2A). We trained animals to fixate a red square at the center of the screen. Following a 200 ms delay, we showed 4 texture images (3 distractors, 1 target, detailed below), arranged in a square. Animals had 1200 ms to register a choice, which we marked as their first fixation of 400 ms or longer on one of the images.

**Figure 2:**
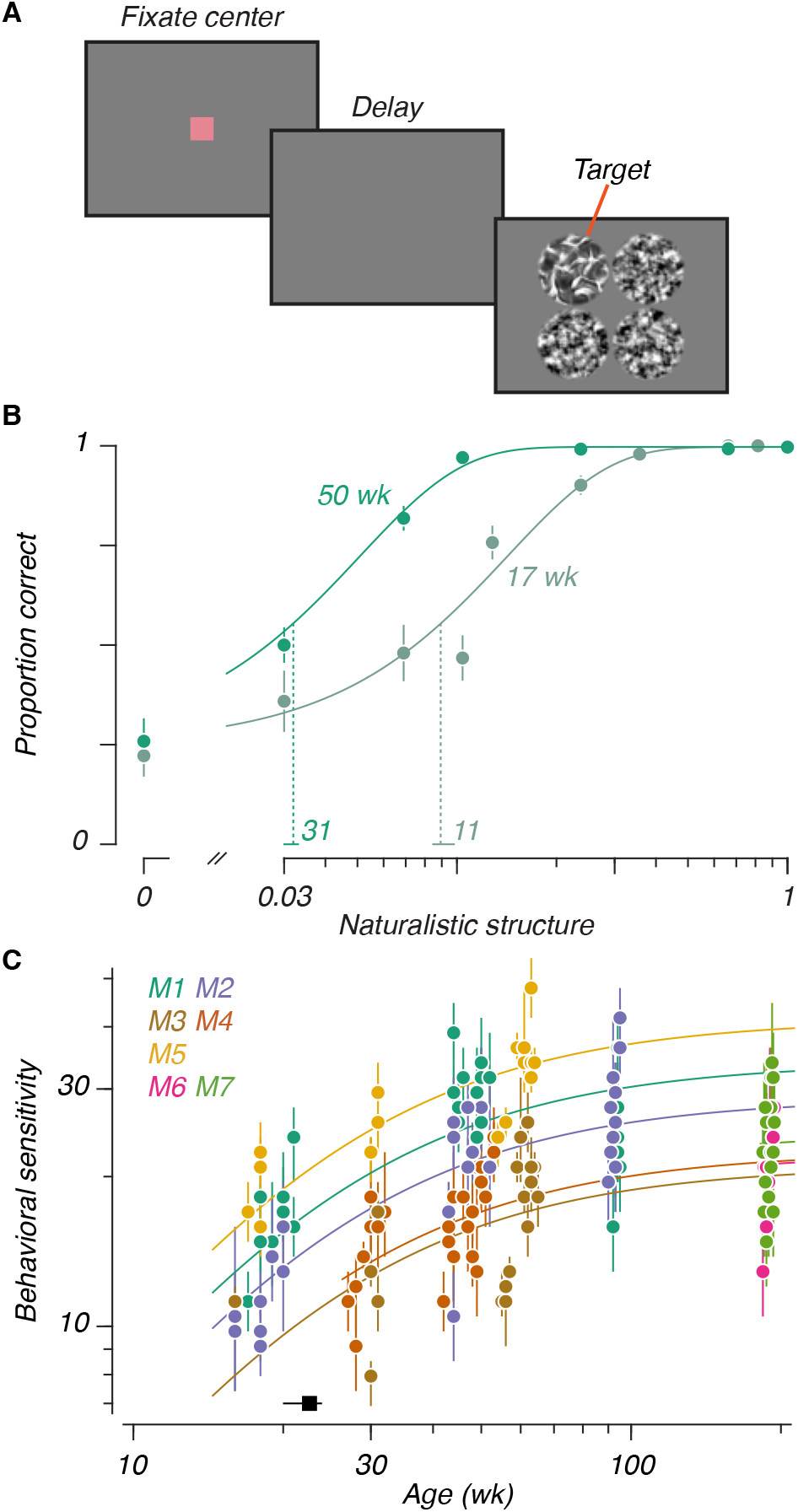
Behavioral task and developmental time course. A: Task design. After fixating the center of the screen, and a 200 ms delay, 4 texture stimuli would appear, each matched in “family” and differing in “sample”. The 3 distractors were noise textures. The target varied in the strength of its naturalistic structure. The display remained on for 1200 ms, unless the animals correctly chose the target by looking at it for at least 400 ms. B: Psychometric functions for one animal at two ages. Filled circles represent data ( binomial variance), Solid lines represent cumulative Weibull psychometric functions fit to the data. Vertical dashed lines mark thresholds, horizontal lines at the base represent 95% CIs. Numerical values above the abscissa reflect sensitivities (inverse thresholds). C: Sensitivities versus age: Each animal is represented with a distinct color. Points represent sensitivities (*±*95% CI), solid lines represent Michaelis-Menten functions corresponding to each animal. The black point above the abscissa represents the half-maximum age extracted from the Michaelis-Menten fit (*±* 95% CI).

We generated all textures using the Portilla and Simoncelli model (Freeman et al., 2013; Portilla and Simoncelli, 2000), including the “noise textures” we used as distractors, which matched the local spectral content of a given natural image. We also generated “naturalistic textures” using the model – textures that additionally matched the local correlation structure of the original image (Figure 1A). By interpolating the model parameters between matched noise and naturalistic textures, we generated textures parametrically varying in the strength of their structure (also see Freeman et al. (2013)). We used one of these naturalistic textures, varying in strength, as the target for each trial. Within a given experimental session, all textures belonged to the same “family” – they were derived from a single ancestral natural image. Within a trial, all 4 images (including distractors) came from different “samples” – no 2 images in a trial were identical. *As a result, the only informative difference between the images was the presence of naturalistic structure in the target*.

Animals learned the fundamentals of the task within a single 30-60 minute session: they could reliably recognize textures at the earliest ages we studied. Their sensitivities stabilized within 3-4 sessions. Figure 2B shows psychometric functions measured from one animal (M1) at 17 and 50 weeks. Performance varied lawfully with the strength of the naturalistic structure at both ages. While performance on highly naturalistic textures was perfect at both ages, performance close to threshold improved with age – sensitivities (vertical dashed lines, numbers listed above the abscissa) more than doubled in this example.

From our sample of 7 animals, we measured sensitivities to 5 different texture families from as early as 16 weeks to as late as 194 weeks (Figure 2C). We modeled the relationship between sensitivity and age with a modified Michaelis-Menten function which had the same shape for all animals. Animals differed systematically in their sensitivity, so we fit a maximum separately for each animal – this model to fit our data the most parsimoniously. We computed a half-maximum age of 23 weeks (50% CI: [20, 24]), suggesting that naturalistic texture sensitivity matured at a rate similar to other spatial vision tasks (El-Shamayleh et al., 2010; Kiorpes and Bassin, 2003; Rodríguez Deliz et al., 2024). A goal of these experiments was to establish a benchmark for comparison with separate neural measurements (below), made across a different age span. We parametrically estimated the magnitude of behavioral change between 26 and 52 weeks. Across this span, we found that sensitivities increased by a factor of 1.39 (95% CI: [1.35, 1.44]).

To confirm that our results did not reflect the learning of particular local features, we introduced a fifth texture family only once animals approached 1 year of age (data are plotted separately for each family in Supplemental Figure 1). Performance in this held-out family was consistent with performance on the other 4, suggesting that sensitivity was primarily determined by age, not stimulus-specific experience. Separately, we tested 2 animals cross-sectionally (M6 and M7), when they were nearly 4 years old (an age when visual capabilities are mature in the macaque). Their sensitivities were consistent with those seen in the other animals at ages at or beyond 1 year, despite their lack of previous experience with this task.

### Single site texture selectivity is stable across development

Having observed that behavioral texture sensitivities increased from 26 to 52 weeks of age, we asked whether neural correlates of this improvement would be visible in the ventral visual areas modulated by the same statistics in adults, including areas V2, V4, and IT, as well as V1, an area previously shown to be insensitive to naturalistic structure (Freeman et al., 2013).

To test this, we recorded multiunit neural activity from 6 passively-fixating animals (see Table 1). We implanted 96 electrode “Utah” recording arrays in V1, V2, V4, and IT. We also implanted an array in the foveal confluence of visual cortex (hereafter FC, after Brewer et al., 2002), for which the specific area could not be determined. In our first experiment, we recorded responses to a total of 35 texture families (Figure 1B), with 15 samples per family, positioned to cover the receptive fields of the sites on the array. Most of our recordings were longitudinal – we recorded in V1, V2, and V4 from 30 to 56 weeks, and in FC and IT from 20 to 36 weeks. We also recorded single sessions at 409 weeks in V4, and 221 weeks in IT.

**Table 1:** Experimental details. Animal numbers reflect the numbers used in behavioral experiments (female: f, male: m). Numbers given in the fourth column represent the number of visually responsive sites used at a given age, for the areas we recorded in that animal, using the texture set of 35 image families. For M3, we were not able to confidently delineate area boundaries in the foveal confluence. Recordings from these arrays may therefore reflect activity in either V1, V2, or V4, and are listed as F.C. in the above table. M6 was implanted with arrays in both hemispheres at different ages, and is listed twice, corresponding to the separate arrays. For the comparison in Figure 3D, we used data recorded at a single age, marked here with an obelisk (*†*).

| Subject | Behavioral age range (wk) | Areas recorded | Recording ages (wk; number of sites) |
| --- | --- | --- | --- |
| M1 (f) | 17-95 | V1-V2 border, V4 | 29 (n = 56, 34, 90), 37<sup>†</sup> (n = 60, 33, 91),<br>56 (n = 61, 32, 95) |
| M2 (f) | 16-95 | V2, V4 | 29 (n = 92, 21), 37<sup>†</sup> (n = 85, 45),<br>56 (n = 92, 57) |
| M3 (m) | 16-65 | FC, IT | 20<sup>†</sup> (n = 87, 39), 31 (n = 81, 40),<br>34 (n = 55, 0 [F.C. only]) |
| M4 (f) | 27-53 | N/A | N/A |
| M5 (m) | 17-64 | N/A | N/A |
| M6 (m) | 184-194 | V1, IT | 30 (n = 0, 30), 34<sup>†</sup> (n = 13, 56) |
| M6 | (see above) | IT | 221<sup>†</sup> (n = 52) |
| M7 (m) | 184-194 | IT | 28 (n = 31), 34<sup>†</sup> (n = 48) |
| M8 (m) | N/A | V4 | 409<sup>†</sup> (n = 96) |

Poststimulus time histograms from all areas are shown in Figure 3A, for sample sites recorded at the youngest (top row) and oldest (bottom row) ages measured in a given area. Texture selectivity – meaning a preference for naturalistic over noise textures – increased successively from V1 to V2 to V4, and declined from V4 to IT. We quantified selectivity in units of d’, and measured selectivities from populations at the youngest at oldest ages in our sample, in response to all 35 texture families (Figure 3B). Selectivity again increased along the ventral stream – measurements in V2, V4, and IT were visibly more selective for texture than V1 or FC. As in Freeman et al. (2013), we observed similar tuning for texture families between sites in a given area – the horizontal bands visible in Figure 3B suggested similar tuning across sites. Qualitatively, we saw evidence of texture modulation from the earliest ages tested in all areas – as early as 20 weeks in IT. One notable feature of these data is that the response dynamics are similar across age for all areas *except* IT, where responses were quite sluggish at 20 weeks. We return to this issue below.

**Figure 3:**
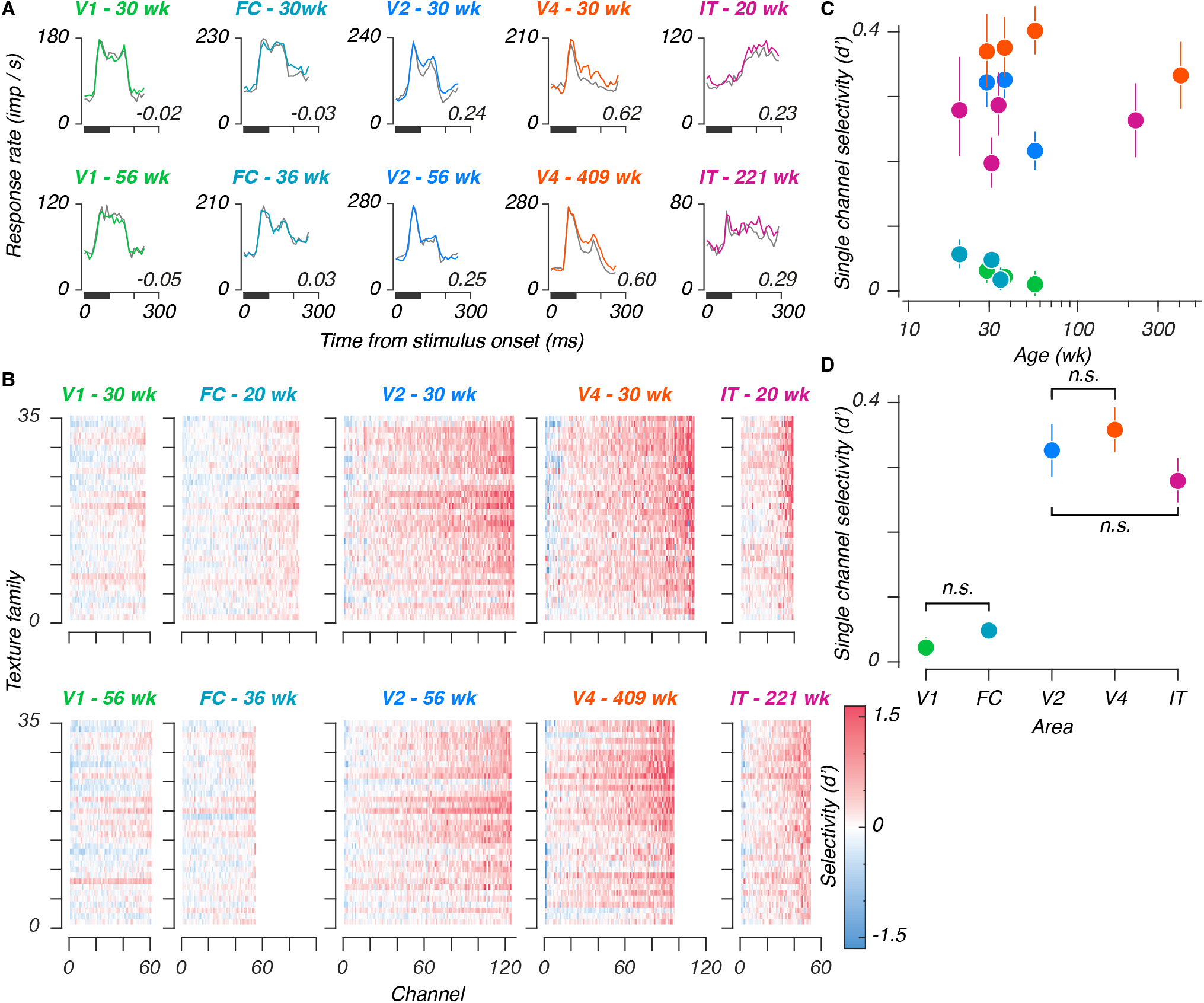
Single site texture selectivities across ages and areas. A: Poststimulus time histograms for example sites, recorded from each area (V1, foveal confluence, V2, V4, and IT), at early and late ages. Traces represent mean responses to all naturalistic textures (color), and all noise textures (gray), for stimuli presented in the first position of a stimulus block, numerical values represent selectivities, for naturalistic versus noise textures, measured across all stimuli. B: Single-site selectivities across all stimuli and areas. Within each panel, each element represents the selectivity (d’) for a given texture family for a given site. Rows are organized from bottom to top based on increasing perceptual sensitivities as reported in Freeman et al. (2013) (their Figure 7). Each column represents 1 site from the total sample measured at a given age, sorted in order of increasing selectivity. The top row of panels represent the youngest age measured in a given area, the bottom row represents the oldest age. Colors were cut off at 1.6, corresponding to 90% of a standard normal distribution. C: Single site selectivities versus age. Points represent mean values, error bars represent 95% CIs. D: Single site selectivities between areas (one age used per array - see Table 1). Conventions as in C. Pairwise interactions were significant unless marked with “n.s.”

We measured the mean selectivity across texture families for each visually responsive site, from all ages and areas (Figure 3C, see also Supplemental Figure 2). We estimated the relationship between texture selectivity and age by linear regression in all areas except FC (where we lacked an older age point). We saw no evidence of age-related improvement. In V1, V4, and IT, slopes straddled zero (V1: -0.07, 95%CI [-0.16, 0.03]; V4: -0.04, [-0.1, 0.02]; and IT: 0, [-0.07, 0.08]) – evidence that texture selective mechanisms were mature from the earliest ages we measured in those areas.

In V2, this slope was negative (-0.4, [-0.56, -0.23]). Rather than reflecting an effect of age, we suspected that this might instead reflect a change in the tuning in our arrays – among the areas selectively responsive to naturalistic textures, V2 was the only area for which we lacked an older, cross-sectional sample. As such, we wondered whether V2 might have been more vulnerable to changes reflecting the ages of the arrays, rather than the animals. In particular, we wondered whether tuning in our V2 arrays may have become more dispersed with time. To measure this, we computed the ratio of variance across texture families to the overall stimulus-driven variance (Supplemental Figure 3). This measure reflects tuning for specific stimulus classes, relative to the spread across all stimuli. A decline in this value would therefore reflect a shift towards signals which, while still visually evoked, may have become less reliably tuned. This ratio remained stable for recordings made in V1 and V4, but declined in V2, confirming that tuning, measured in this manner, may have declined with age in V2.

### Single site texture selectivity is lower in IT than in V4

We wanted to compare texture selectivities between areas, to compare our results with prior measurements in V1 and V2, to relate the selectivities of V2 and V4 using the same stimuli, and to furnish a first measurement of naturalistic texture selectivity in IT. To avoid double-counting, we chose one recording per array (Figure 3D, and see Table 1). We found a significant group-level difference between areas (F = 14.3, permutation ANOVA p < 10^-5^), and significant pairwise differences between V1 and each of V2, V4, and IT (mean d’ difference: 0.31, 0.34, 0.27, respectively, permutation test p < 10^-5^).

Downstream from V1, we found that selectivity was similar between V2 and V4 (mean difference: 0.03, p = 0.28), in line with prior findings (Okazawa et al., 2017). Selectivities were significantly lower in area IT than in V4 (mean difference: -0.08, p < 0.002), but did not differ significantly from V2 (mean difference: -0.05, p = 0.09). These results suggest that selectivity to naturalistic texture may peak in V4.

### Neural populations stably encode naturalistic textures across development

Our analysis of naturalistic texture encoding at the level of single electrode sites revealed two main findings: first, that texture selectivities were mature as early as could be measured – even as behavioral sensitivities continued to mature. Second, we found that texture selectivities peaked in midlevel visual areas, and declined in IT. We wondered whether these observations also held at the population level. To address the first point, we asked whether population representations of texture – taken across sites – might reveal a link to behavioral development.

To measure how neural populations encoded naturalistic texture, we projected the responses into a high dimensional space in which each axis represents the response of one site, and found the hyperplane that best separated responses to naturalistic and noise textures for each texture family. We then measured the distance between held-out examples of the two texture types along the line orthogonal to the hyperplane as a d’. As an analogous measure of *detectability,* we constructed identical decoders measuring the distance between naturalistic (or noise) textures and blank stimuli.

Performance values for our naturalistic texture discrimination scheme are plotted in Figure 4A, for populations of 30 sites (a number chosen to facilitate comparisons – see Methods). In all areas, the encoding of naturalistic textures was stable from the earliest ages measured, including both longitudinal measurements made within the first year of life (as early as 20 weeks), and, in V4 and IT, recordings made at considerably older ages (roughly 8 and 4 years, respectively). We fit slopes relating population performance with (log-transformed) age, and used those slopes to estimate the magnitude of change between 26 and 52 weeks (recall that behavioral sensitivities increased by a factor of 1.4 over this time). As for single sites, changes in behavior were not accounted for by changes at the population level – in V4 and IT, performance at 52 weeks fell to 0.95 and 0.97 of the performance at 26 weeks (95% CI: [0.93, 0.97], [0.96, 0.98], respectively). In V2, we saw a substantial decline, with performance falling to 0.69 of the 26 week estimate (CI: 0.64, 0.74), which can be attributed to the inferred change in recording quality discussed above. V1 was more volatile, reflecting its overall lack of sensitivity to naturalistic texture (Change between 26-52 weeks [CI]: 2.5 [2.1, 2.9]). We did not consider FC for this analysis, as we did not sample it at later ages.

**Figure 4:**
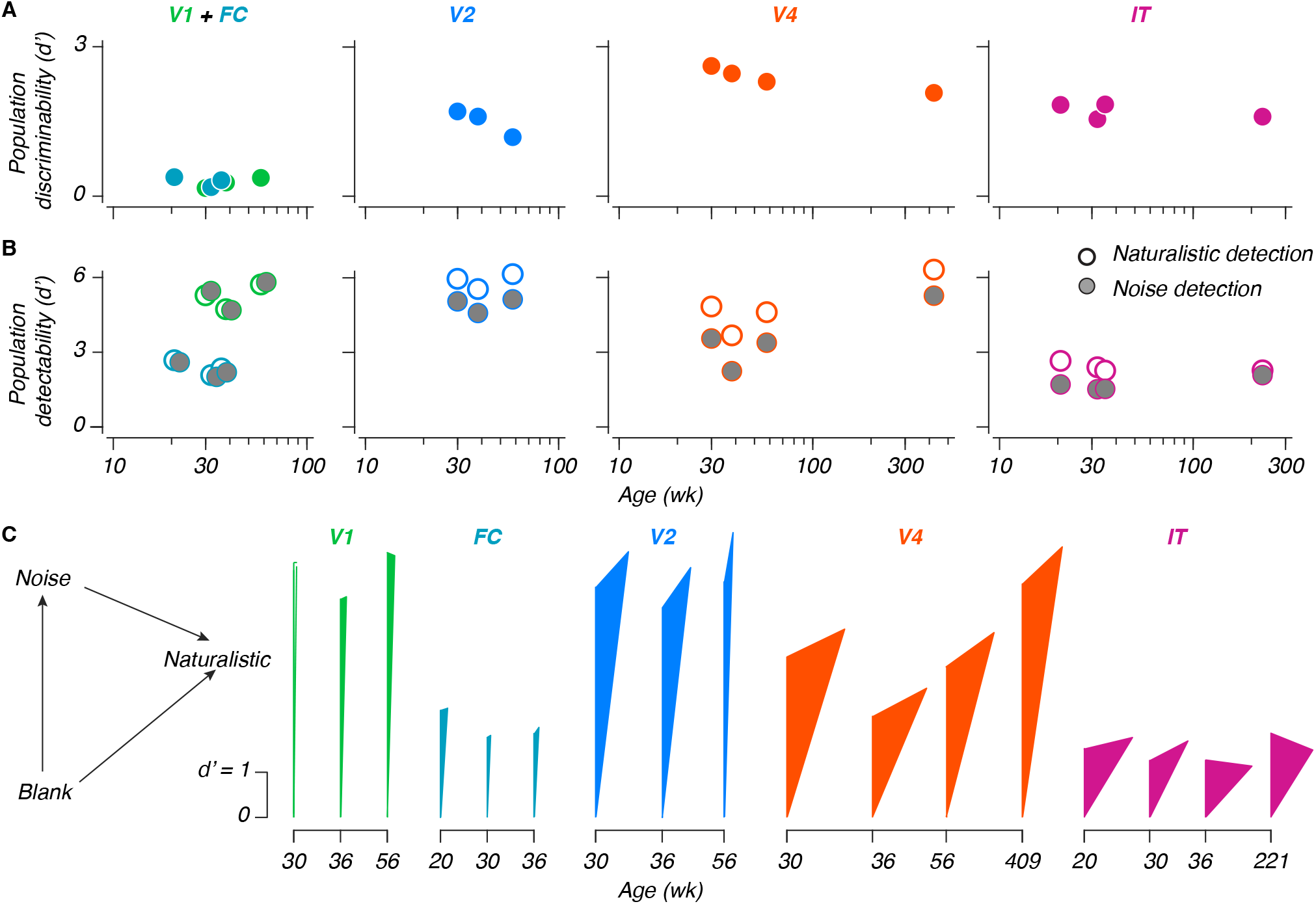
Population coding and representations in the developing ventral stream. A: Naturalistic texture discriminability between naturalistic and noise textures, versus age. Values are the mean across multiple population samplings of 30 sites. B: Stimulus detection performance versus age, for either naturalistic textures (white points), or noise textures (gray points, shifted horizontally when necessary for visibility). Note that the scale on the ordinate is halved in height relative to A; conventions are otherwise the same. C: Triangular representation. As indicated in the legend on the left, the length of each side reflects the 3 population metrics from panels A and B. The sides emanating from the bottom reflect detectability; the far side reflects discriminability. NB: The leftmost measurement in V1 could not geometrically form a triangle – it has been left hollow to reflect this.

Our measurements of texture detectability provided additional context (Figure 4B) – detectability remained stable for both texture types during development, but was greater than texture discriminability (reflecting the ability of naturalistic textures to reliably evoke visual responses). Using the same 26-52 week comparison, we found modest increases in V1 and V4 (112% [CI: 107,117], 113% [112, 115], respectively), a modest decrease in IT (97% [95, 99]), and no significant change in V2 (105% [99.8, 110]). We again did not consider FC in this analysis. Together with our single site measurements, these results suggested that the increase in behavioral sensitivity to naturalistic textures that we observed did not stem from developmental changes in the ventral stream, and must instead result from changes elsewhere in the brain.

### Population representation of texture in different areas

We recorded population responses to naturalistic texture in V1, V2, V4, and IT. We wondered how well the activity in each area could support texture discrimination. We compared the performance of each area using our measurements of discriminability (Figure 4A, see also Supplemental Figure 4). Performance was poor in V1 and FC, higher in V2, and highest in V4. IT performance was lower, similar to V2.

Comparing texture discriminability in different areas revealed that these values did not always reflect the simple difference between the *detectability* values for naturalistic and noise textures. An additive relationship between detection and discrimination would imply that the population representation of naturalistic structure is in the same subspace as the one that represents the presence of a texture stimulus. Deviations from an additive relationship indicate that the population encoding of naturalistic structure uses a different population subspace.

To measure the relationship between the population representations for naturalistic structure and texture detection, we draw triangles whose side lengths reflect the 3 population measurements from each neural sample – the detectability measurements for noise and naturalistic texture, and the measurement of discriminability between the two. Figure 4C depicts these triangles. The length of the left, vertical arm represents an area’s ability to detect noise textures. The length of the right arm (arising from the bottom) represents its ability to detect naturalistic textures. Finally, the length of the opposite arm corresponds to the discriminability between naturalistic texture and noise.

Neurons in V1 and V2 respond similarly strongly to the presence of texture images. As a result, the heights of the triangles in V1 and V2 are comparable. On the other hand, neurons in V2 respond *differentially* to naturalistic versus noise textures. As a result, the opposite side in V2 is longer than that in V1 (where the opposite side is of negligible length). Critically, the opposite side is not only *longer* in V4, the triangles formed are *wider,* suggesting that naturalistic structure is not only more strongly represented in V4, it is represented in a population subspace increasingly different from the one corresponding to texture detection.

In IT, the *height* of the triangles is less than that in V1, V2, or V4, whereas their relative width is similar to those of V4, suggesting that the change in the subspace of the naturalistic texture representation in V4 may be preserved in IT. These results reflect a transition from the detection-driven response of V1, towards a representation in V2 and V4 which is both increasingly sensitive to naturalistic structure, and which represents this structure along a subspace increasingly different from the one corresponding to texture detection. In IT, while texture discriminability was lower than in V4, the dimensionality of how naturalistic texture is represented was similar.

### IT dynamics become faster during development

Visual response latencies in areas V1 and V2 shorten during the first 8-16 weeks of life (Rust et al., 2002; Zheng et al., 2007), and latencies in area IT are longer in infancy than in adulthood (Rodman et al., 1993). Because latency shifts can reflect morphological changes like myelination, we wondered whether latency measurements might reveal visual development. To study changes in dynamics, we measured population performance using the same texture discrimination and naturalistic texture detection schemes as before, training and testing performance separately for the spike count in each 10 ms time bin in our data, as opposed to the 150 ms window used in the above analyses (see Hung et al. (2005), their Figure 3B).

We used these curves (Figure 5A) to investigate whether neural dynamics changed developmentally. We measured the onset latency as the time at which a performance curve first began to rise from baseline (in V1 and FC, we only measured latencies in the detection paradigm). Latencies remained largely stable for V1, V2, and V4, for both paradigms (Figure 5B). Latencies in IT shortened during the measured span. While saturation in areas V1 and V2 is unsurprising, given prior reports (Rust et al., 2002; Zheng et al., 2007), previous recordings from infant IT were made at the age of 4 weeks (Rodman et al., 1993) – our measurements extended that maturation out to 20 weeks. Separately, comparing our simultaneous measurements from V2 and V4, we observed that naturalistic textures were discriminable in V2 at earlier times than in V4, despite the latter area’s overall heightened discriminability, evidence that this signal may first emerge in V2.

**Figure 5:**
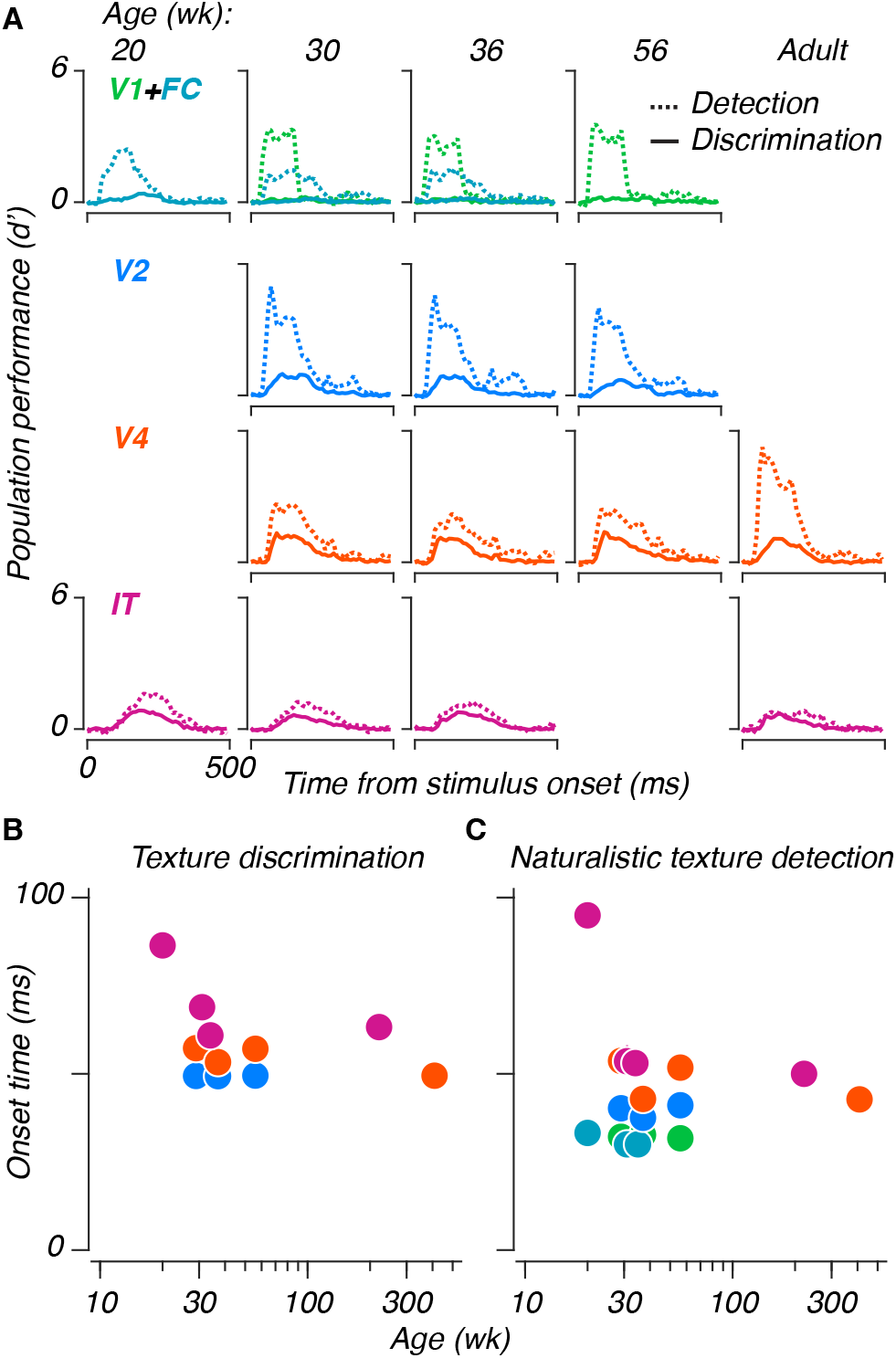
Age-related changes in the temporal dynamics of responses in IT. A: Population performance curves versus time, trained and tested separately for each 10 ms time bin. Lines represent means across population samples. Dashed lines represent naturalistic texture detection, solid lines represent texture discrimination. B: Half maximum latencies, measured as the first time at which performance exceeded half of the eventual maximum, for both paradigms. Error bars represent 95% CIs across population resamples; they are often smaller than the symbols, and thus invisible.

### Population neurometric sensitivities are stable across development

*Behaviorally,* even young animals were largely perfect at discriminating fully naturalistic textures from noise (e.g. Figure 2B) – developmental improvement was only noticeable near threshold. The neural measurements we reported above sampled a large number of texture families, but included only fully naturalistic and noise textures. To test for evidence of neural development near threshold, we recorded neural responses to the texture images we used in our behavioral experiments – textures varying in the strength of their naturalistic structure. We recorded these responses from 3 of the animals reported above, again during passive fixation, and measured neurometric performance from populations of 20 sites.

To measure neurometric sensitivities, we again found the hyperplane best separating fully naturalistic and noise textures. To simulate performance on a 4-alternative task, we projected 4 individual neural responses onto this axis – 3 noise texture responses (simulated distractors), and 1 trial varying in naturalistic structure (simulated target). If the magnitude of the target projection was the largest, we scored the trial as correct. Otherwise, we scored the trial as incorrect. We iterated across all responses in our held-out test set, extracted proportions correct for each level, and fit neurometric sensitivities. Examples are depicted in Figure 6A. The relationship between areas at the level of sensitivities was similar to what we previously observed with discriminabilities – V1 and FC were weakly sensitive to naturalistic textures, if at all. Sensitivities grew moving from V2 to V4, and declined moving from V4 to IT.

**Figure 6:**
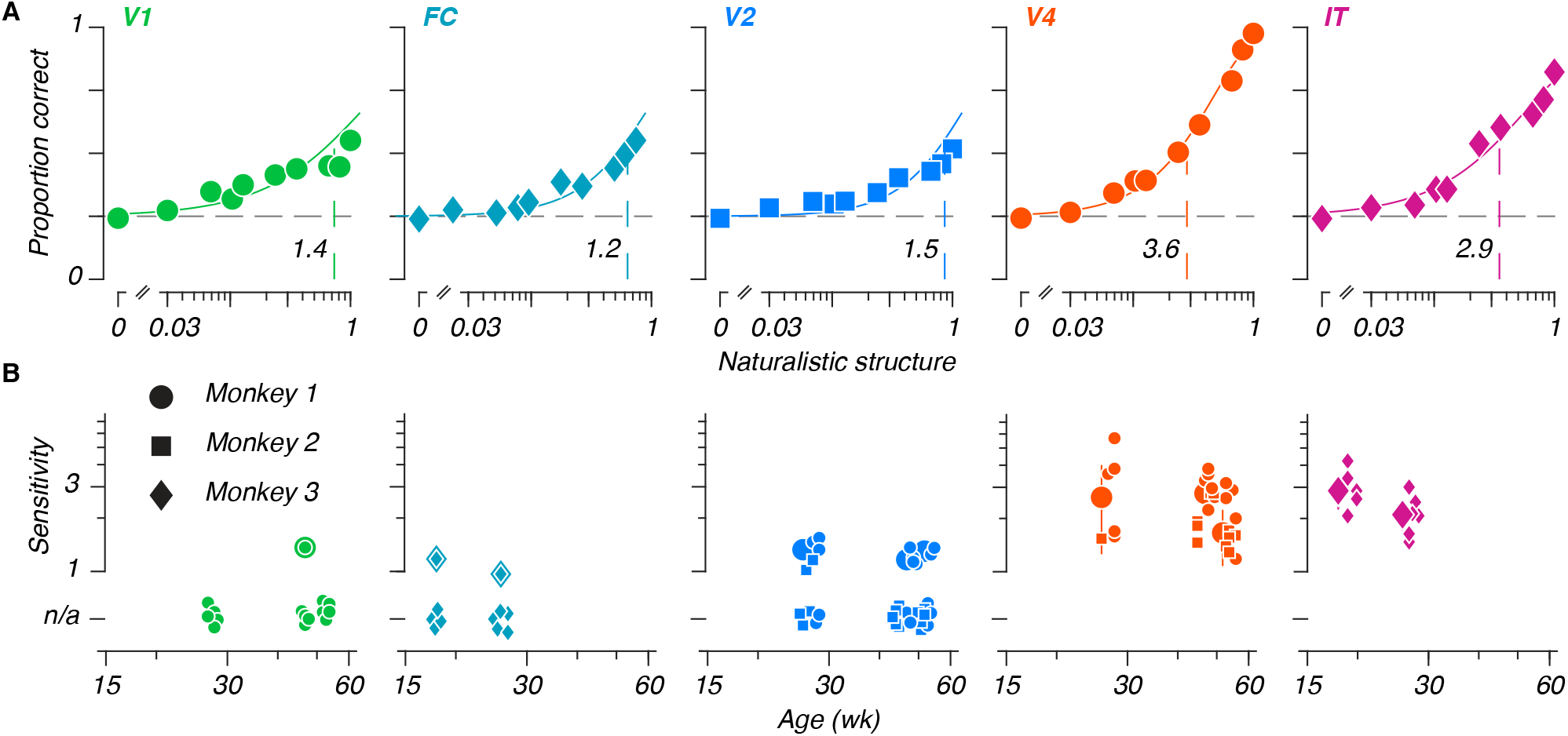
Stable neurometric sensitivities in the developing ventral stream. A: Example neurometric functions from each area. Each panel depicts a neurometric function, and the population average for each condition, for one example session. Points represent the average proportion correct for a given level, solid lines represent neurometric functions fit to the data. Vertical dashed lines represent thresholds, which were inverted to obtain sensitivities (black numeric values). B: Neurometric sensitivities versus age. Large points represent median sensitivities across 20 site subsampled populations, for all fitted data measured in a given area, at a given age. Small points represent the sensitivities measured for individual sessions (symbols indicate animal identity). Sessions for which a sensitivity could not be extracted are represented with points below the ordinate (”n.s.”). Error bars on large points depict median absolute deviations across fit sensitivities.

We extracted sensitivities from all sessions (Figure 6B). Our observations were largely consistent with our previous reports – sensitivities increased in downstream areas, and did not change as a function of age. While neural sensitivities seem to be lower than behavioral ones, this merely reflects the chosen population size (see Supplemental Figure 4). We repeated these measurements after shuffling the relationship between sites, and found a modest decrease in performance that did not vary with age. Finally, we measured discriminability between fully naturalistic and noise textures in these same data, to relate discriminability with sensitivity at threshold (Supplemental Figure 5), and found that values tracked closely. Between population sensitivities, discriminabilities, and the correlation structure between sites, naturalistic texture encoding in the ventral stream was stable across all ages tested, despite an increase in behavioral sensitivity during the same span.

## Discussion

### Visual development

We measured the rate at which macaque monkeys improved in their behavioral sensitivity to naturalistic textures. Sensitivities increased during early life at a rate similar to other spatial vision tasks (El-Shamayleh et al., 2010; Kiorpes et al., 2012; Rodríguez Deliz et al., 2024; Stavros and Kiorpes, 2008). Based on these results, and given prior evidence that neurons in areas V2 and V4 are sensitive to the same naturalistic structure, we recorded multiunit neural sensitivities in areas V1, V2, V4, and IT to naturalistic textures, using a superset of the images used for behavioral measurements. Other than changes in response dynamics in area IT, we saw no evidence for neural development of the representation of naturalistic texture in any of the ventral visual areas from which we recorded.

Previous measurements from early visual areas established that both tuning and timing in areas like the LGN, V1, and V2 reach maturity by or before 16 weeks (Movshon et al., 2005; Rust et al., 2002; Zheng et al., 2007). Previous measurements from IT suggest that neuronal responses are selective for object identity from as early as 4 weeks (Rodman et al., 1993), and that response onsets are immature relative to adults. Through the use of population analysis methods, and our use of a behavioral benchmark, we have extended the disconnect between behavioral and neural development to the end of the ventral visual stream.

The stable neuronal representations we observed across development complement recent results obtained from infant humans and macaques using functional imaging. In both species, retinotopic organization has been observed across a number of ventral areas (including V1, V2, and V4), from early ages – including the first 1-2 weeks in macaques (Arcaro and Livingstone, 2017), and 5.5 months in humans (Ellis et al., 2021). In infant macaques, face-preferring patches in IT are visible from the first month of life (Livingstone et al., 2017). In infant humans, face patches can be seen in IT-analogous areas from the age of 4 months (Deen et al., 2017). While refinements of these selectivities have been observed, it is of note that that refinement lags the onset of face-guided behavior in infants of both species (Maurer and Salapatek, 1976; Mendelson et al., 1982). The observed changes may instead reflect a process distinct from feed-forward neuronal drive. Taken with our own findings, evidence suggests that the visual system is both anatomically and functionally mature from an early age; behavioral development may therefore reflect development downstream from ventral visual cortex.

If behavioral development is not limited by sensory cortex, what might constrain it? One possibility relates to decision and action – the lateral intraparietal area (LIP), frontal eye fields (FEF), and superior colliculus (SC) represent information pertaining not just to the stimulus, but also its relation to an eventual behavioral choice in saccade-based tasks (Gold and Shadlen, 2007; Mirpour and Bisley, 2021). Whereas neuronal motion sensitivity in macaque LIP correlates with behavioral improvements in a perceptual learning paradigm, the sensitivity of MT neurons remains stable (Law and Gold, 2008). We have previously shown a relationship between the visual stimulus used in a task, and the corresponding rate of behavioral development (see Kiorpes, 2015). In particular, behavioral development tends to take longer for tasks which require the integration of stimulus information across relatively large spatial expanses. As a result, performance may be more dependent on development in downstream areas with large receptive fields, such as association areas related to visual recognition, which may develop more slowly. Protracted development in downstream areas is supported by anatomical measurements – occipital areas mature more quickly than parietal and frontal areas in macaques (Scott et al., 2016) and humans (Huttenlocher and Dabholkar, 1997). We designed our task to obtain psychophysical data as quickly as possible. A task with a more explicit delay period could allow for similar measurements in LIP, FEF, or any other area spanning the space between the ventral stream and motor output, assuming that such a task could be learned by infants.

More broadly, these results, along with other demonstrations of adult-like tuning in the developing primate visual system, may help reorient our thinking about visual development (see also Makin and Krakauer (2023), for a contemporary reevaluation of postnatal reorganization). Earlier studies of plasticity following monocular deprivation led to a focus on the malleability of the developing visual system (Blakemore and Van Sluyters, 1974; Carlson et al., 1986; Movshon and Blakemore, 1974; Wiesel and Hubel, 1963), as opposed to studies that demonstrated the relative maturity of the normal visual system (Hubel and Wiesel, 1963). Moving forward, it may be worth thinking of visual cortex as a series of areas which are mostly adult-like in their tuning and function during typical development, and which feed into downstream areas whose ability to effectively process those sensory inputs may develop more slowly.

### Visual processing

While the primary focus of these experiments was to understand development, our measurements also enabled us to explore naturalistic texture encoding in the areas from which we recorded. Like others, we found selectivity for texture information in mid-level visual areas like V2 and V4 (Arcizet et al., 2008; Freeman et al., 2013; Kim et al., 2019, 2022; Okazawa et al., 2017; Yu et al., 2015; Ziemba et al., 2016). Our latency measurements suggest that neural signals supporting texture discrimination emerge first in V2, and are amplified in V4. Our population analyses found that naturalistic texture was more strongly represented in V4 than in V2, despite similar single site metrics. We also found that the neural population subspaces supporting stimulus detection and naturalistic texture discrimination were more different in V4 than in V2.

Our measurements from area IT showed lower responses to textures than in earlier areas. In our data, naturalistic texture information is most strongly represented in V4. Yet despite the diminished performance in IT, both areas appear to use distinct neural subspaces for naturalistic texture discrimination and for stimulus detection. An attenuation of texture responses in IT is consistent with its known sensitivity to visual objects (Gross et al., 1972; Hung et al., 2005; Majaj et al., 2015; Rust and DiCarlo, 2010; Tanaka, 1996). In that light, our results suggest that a neural representation of naturalistic structure first emerges in V2. Those signals are amplified in V4, and are represented in a subspace increasingly different from that reflecting the simple presence of the stimulus. In IT, the response to textures is attenuated, but the dimensionality is preserved. Combined with separate V4 representations of shape (Kim et al., 2019) and color (Bushnell et al., 2011), the neural subspaces we observed may therefore reflect a coding strategy in IT which can support scene segmentation and object-centric coding while still providing information about more elementary visual features (DiCarlo et al., 2012; Lettvin, 1976; Movshon and Simoncelli, 2014).

## Acknowledgements

We are grateful to members of the Visual Neuroscience Laboratory for advice and discussion; Mike Hawken, for comments on the manuscript; Dan Sanes and Jonathan Levitt for discussions on the results; and to Michelle Hernandez, Tiffany Tang, and Kahlia Gronthos for help with data collection. This work was supported by grants from the National Institutes of Health – including R01-EY024914, R01-EY031446, F31-EY031249 (GML), T90-DA043219 (GML), T32-EY007136 (JAM, GML), T32-MH019524 (LK, CLRD) F31-EY031592 (CLRD), and F31-EY026791 (BNB).

## Materials and Methods

### Visual stimuli

We generated stimuli using the texture model of Portilla and Simoncelli (2000), using the methods detailed in Freeman et al. (2013). For our measurements of behavioral and neural texture sensitivities, we used 5 texture image sets, each corresponding to one texture “family” based on a single ancestral natural image (Figure 1A). As in Freeman et al. (2013), the resulting textures varied in the strength of naturalistic structure and in the precise location of their elements (their “sample”). We measured responses to one texture family per session. For behavioral measurements, we used 15 samples for each level of naturalistic structure from a given family; for neural measurements we used 5 samples. In both cases, the strength of structure varied from 0 (spectrally matched noise) to 1 (fully naturalistic).

For a separate measurement of naturalistic texture discriminability, we used a larger texture image set of 1150 images (Figure 1B): 525 naturalistic and noise textures (1050 total, from a total of 35 texture families, and with 15 texture samples per family), and 100 blank images (used to measure baseline firing rates).

We presented stimuli on a gamma-corrected CRT monitor with a mean luminance of 28 cd m^-2^, a resolution of 1280 by 960 pixels, and a frame rate of 100 Hz. We seated animals in a custom primate chair 114 cm from the monitor, at which the monitor subtended 20 by 15 deg.

### Behavioral experiments

We performed all animal procedures in accordance with the National Institutes of Health *Guide for the Care and Use of Laboratory Animals* (2011), and with the approval of the New York University Animal Welfare Committee.

We trained 7 macaques on our behavioral task (*Macaca nemestrina,* 3 female), using standard operant conditioning methods. Animals initiated trials by fixating a 1-2 deg red square at the center of the screen. Once they had done so, the square disappeared, and the screen remained blank for 200 ms. Four texture stimuli then appeared – 3 noise texture distractors, and 1 target texture. Each texture was 6.4 deg in diameter, centered *±*3.2 deg from the center of the screen in both the horizontal and vertical direction (thus 4.5 deg eccentric). Once the textures appeared, subjects had 1200 ms to register a choice, which was defined as fixation on 1 of the 4 stimuli for more than 400 ms. Trials ended immediately after correct responses, at which point subjects received a juice reward. There was no penalty for incorrect responses, except that the stimuli would remain on screen for the full 1200 ms. We did not analyze trials in which the animal failed to respond, which were rare.

We measured naturalistic texture sensitivity using a single texture family per session. On each trial, we varied the position of the target among the 4 locations as well as the strength of its naturalistic structure. The naturalistic structure varied along a series of fixed levels. We used different samples for all 4 images – including the noise texture distractors, to ensure that subjects could only distinguish targets from distractors on the basis of naturalistic texture statistics, instead of pixel-level cues.

We used the method of constant stimuli to determine the difficulty level for each trial. Roughly 5% of trials were catch trials in which the target was also a noise texture – we rewarded animals for choosing this target in the usual way. The levels we chose for a given session were adapted to span the psychometric function from chance to perfect performance, while maintaining the animals’ motivation. To maintain overall motivation levels, we showed fully naturalistic textures disproportionately often.

Sessions typically contained 600-1000 total trials, with roughly 100 trials per condition. We measured performance on each texture family multiple times at each age, with the exception of M3, due to constraints imposed by electrophysiological measurements. We measured performance on at least 4 texture families per animal. We introduced a fifth for some animals at older ages, as a way to both estimate animals’ ability to generalize on the task, and to probe whether their performance was learned or age-based. We excluded sessions containing fewer than 120 trials, or with a peak performance of less than 90%.

### Physiological experiments

For physiological recording, we trained the animals to fixate the central 3 deg of the screen, which was marked with a red central square 0.1-0.2 deg across. Texture images appeared after 160 ms in a pseudorandom order. For our texture set with 35 texture families, we presented blocks of 8 images, and showed each image for 100 ms, followed by a 100 ms blank interval. For our measurements of neurometric thresholds, we showed blocks of 4 images for 200 ms (the same used to measure behavioral thresholds), and used a 200 ms blank interval.

All texture images measured 6.4 deg in diameter. For most data, receptive fields (detailed below) were centered within the central 1.5 deg – for these cases, we centered stimuli at the center of the monitor. For one animal, detailed below, receptive fields were roughly 8 deg from the center of gaze. In this case, we centered stimuli over our estimate of the receptive field center. In all cases, the visual stimuli covered the aggregate receptive fields of the recording sites.

We rewarded animals for remaining fixated through a block with a juice reward. If an animal broke fixation during a stimulus, we interrupted presentation and presented the interrupted stimulus again later in the overall sequence. We recorded at least 4 repetitions of each stimulus from our multifamily texture image set. We recorded at least 36 repetitions of each stimulus in our (smaller) image set of textures varying in their naturalistic structure.

### Neural recording

We recorded multiunit neural responses from a total of 6 animals. We recorded longitudinal data from 5 of these animals, starting between 18 and 26 weeks, and continuing for as long as the recording arrays could be maintained (see Table 1). We implanted one of these animals a second time after it had reached 4 years of age. Finally, we recorded from area V4 of a separate adult. We combined data collected from similar ages (within 2 weeks).

After training animals to perform the fixation task, we implanted 96 electrode (Utah) recording arrays under sterile conditions. We used gross anatomical features to inform decisions about array placement. Electrodes were arranged in a 10*×*10 square (four positions were not used for recording), had shanks 1 mm long and had an interelectrode spacing of 0.4 mm. Following implantation, we used anatomical landmarks (including gross anatomical landmarks such as sulci and vasculature, which we observed surgically and related to previously documented area boundaries Saleem and Logothetis (2012); Winters et al. (1969)), physiological response properties (e.g. response latencies), and receptive field characteristics to determine the cortical location of array sites. All sites on an array were typically within a single visual area, with 2 exceptions. In one animal, one array lay on the border between V1 and V2. In the other animal, we were unable to determine whether the arrays were located in area V1, V2, or V4. As receptive fields were close to the center of gaze, we refer to these data as stemming from the “foveal confluence” (see Brewer et al. (2002), their Figure 8B), as opposed to one or another visual cortical area.

We recorded bandpass filtered (250 Hz to 7.5 kHz) electrical activity, at a sampling rate of 30 kHz. To minimize the influence of common-mode signals, we subtracted the median sample-by-sample voltage across all sites, and then defined spike events (threshold crossings) as negative voltage deviations at least 3 times the root mean squared deviation of the baseline voltage.

### Behavioral data analysis

For the 5 animals we tested longitudinally, we used between 22 and 33 total sessions. For the 2 animals we tested at older ages, we used 3 sessions in one, and 11 in the other. For each animal, we combined behavioral data from a given texture family collected within a 7 day span.

Two animals (M3 and M5) acquired a tendency to choose the same spatial position at older ages. Compared to the other animals in our sample, these two had run more 4 choice oddity tasks of various sorts. We retrained them to mitigate this spatial bias, and rejected sessions where they chose one target at least 40% of the time (all results and model predictions were similar with and without this inclusion criterion).

To measure behavioral sensitivities, we fit a shared cumulative Weibull psychometric function to all data (Wichmann and Hill, 2001a). We used a maximum likelihood fitting process to extract a slope parameter, common to all data, and separate thresholds and lapse rates for each individual measurement. We computed thresholds as the intercept of the Weibull function with 55% correct, corresponding to a d’ of 1 (Hacker and Ratcliff, 1979). We then estimated the individual variability for each threshold estimate using a nonparametric bootstrap (Kingdom and Prins, 2016), for which we fixed the slope and lapse rate to the values extracted from the original fitting routine. We inverted thresholds to obtain sensitivities.

We modeled the relationship between sensitivity, *s,* and age, *a,* using a function of the form:

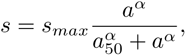

where *s_max_, a*_50_, and *α* are free parameters representing the maximum sensitivity, the age corresponding to half-maximum performance, and a fit exponent capturing the rate of increase (larger values correspond to faster saturation), respectively. We fit this function by minimizing the squared error of the model predictions. Our psychometric function is fit in a logarithmic space (curves sharing a slope and lapse rate, but differing in threshold have the same shape in this space), thus we log-transformed our estimates prior to measuring model error. We tried a variety of models of this general form, and compared them using the corrected Akaike’s Information Criterion (Motulsky and Christopoulos, 2004). Across all such models, our data were best explained by one in which the *a*_50_ and *α* parameters were shared across all data, and the *s_max_* was fit separately for each subject. To estimate the variability of model parameters, we nonparametrically resampled our data, and fit our model to the resampled data. We repeated this 1000 times, and extracted 95% confidence intervals around the parameter estimates.

### Physiological data analysis

#### Initial analysis

For analysis, we binned multiunit threshold crossings into 10 ms windows. To determine single-site response latencies,

we separately measured the discriminability between blank stimuli, and either naturalistic texture or noise texture

stimuli as a d’ between response distributions: 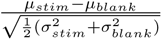, where *µ* reflects the mean of a distribution, and *σ* its standard deviation. We defined the onset time for a given site as the first deviation above a d’ of *±* 0.4, as long as the following 2 windows were also suprathreshold (and of the same sign). The onset time for a given population was the median onset time across visually responsive sites. For most analyses, we then summed responses across the 150 ms window following stimulus onset.

To determine whether a site was visually responsive in a given recording session, we measured the average response separately for even and odd repetitions of each stimulus. We measured the Pearson correlation between these vectors, and used sites with a correlation of 0.4 or greater (the results reported here remained stable across a variety of different metrics and thresholds).

We measured receptive field locations by recording neural responses to a small spot, which was tiled across the visual field. We used d’ values, summed for 200 ms following response onset, between spot stimuli and blanks to estimate receptive field centers and sizes. With one exception, that of M8, receptive fields were located within 2 deg of the center of gaze. For M8, receptive fields were roughly 8 deg eccentric from the center of gaze.

As a single site measure of naturalistic texture discriminability, we measured the d’ between naturalistic and noise texture evoked response distributions, using the above formula.

To compare tuning at the single site level, we measured the variance across all texture families (including naturalistic and noise textures), after averaging across stimulus repetitions, and computed its proportion of the overall stimulus-driven variance.

#### Population analysis

To measure population discriminability, we first performed singular value decomposition on a training set *USV ^T^* = *M_train_* of naturalistic and noise texture responses, for a given population (*M_train_* was a matrix organized in the form stimulus *×* site). Array recordings result in varying population sizes. We primarily addressed this by subsampling populations to a matched size, allowing us to directly compare performance between ages and area. We obtained these subsamples by randomly selecting from the pool of visually responsive sites. We repeated this random sampling to estimate variability across measurements. For our first physiological experiment, using textures from many families, we used populations of 30 sites. For our measurements of population neurometric sensitivities, we used populations of 20 sites. In Supplemental Figure 4, we measured the relationship between population size and performance, and found that our results generalized across population sizes.

To minimize overfitting, we reduced the dimensionality of the resultant basis set *V* to rank 20 (10 for neurometric sensitivities). We then fit a linear discriminant, *b,* to the rotated matrix *M_train_V,* best separating naturalistic and noise responses. We measured performance across training-testing splits by projecting our held out data *M_test_* onto the same axis as *M_test_V b.* We measured the discriminability between naturalistic and noise texture response distributions as a d’. To measure stimulus detectability, we repeated this process, but using naturalistic textures and blank stimuli.

For our first experiment, we measured performance separately for each texture family. For cross-validation, we trained on 14 of the 15 samples, and tested on the held out sample. We reported performance as the average across all 15 cross-validations.

For our analysis of temporal dynamics, we measured discriminability using the same population scheme, but trained and tested separately for each 10 ms time bin. To measure population latencies, we fit a Heaviside function, modified to include a finite slope (which we fit as a free parameter), to the resultant average performance curves.

For our measurements of population neurometric sensitivities, we again used samples for fivefold cross-validation, using 4 samples for training, and the 5th for testing. In addition to population discriminability, we measured proportion correct in analogy to a 4 choice task, by iterating through each response in *M_test_,* with a simulated trial structure, where we projected each individual response in *M_test_* onto our learned axis (simulated target), along with 3 randomly chosen noise texture responses from *M_test_* (simulated distractors). If the projection of the simulated target onto the learned discriminant axis was larger than that of all 3 distractors, we scored the stimulated trial as correct. Otherwise, we scored it as incorrect. After repeating this process for all entries in *M_test_,* we computed an overall proportion correct for each level of naturalistic structure.

We then fit neurometric functions with cumulatives of the Weibull distribution, fit separately to each session where the maximum proportion correct reached at least 0.5. We estimated variability using a parametric bootstrap (Wichmann and Hill, 2001b).

We measured neurometric sensitivities separately using a correlation-based classifier. In this approach, we replaced the projection of 4 trials (a target and 3 distractors) onto a discriminating hyperplane, with the correlation between the same 4 trials, and the average response to fully naturalistic textures, as measured from our training set. Here, we simulated choice as the trial with the highest correlation. Our results in this framework were qualitatively similar to those using the linear discriminant.

For our analysis of the influence of neural correlation structure on decoding, we recomputed the population discrim- inability as measured from our second experiment, after randomly and separately shuffling the trial order for each site, to disrupt any trial-to-trial correlations in neural activity.

---

**Supplementary Figure 1:**
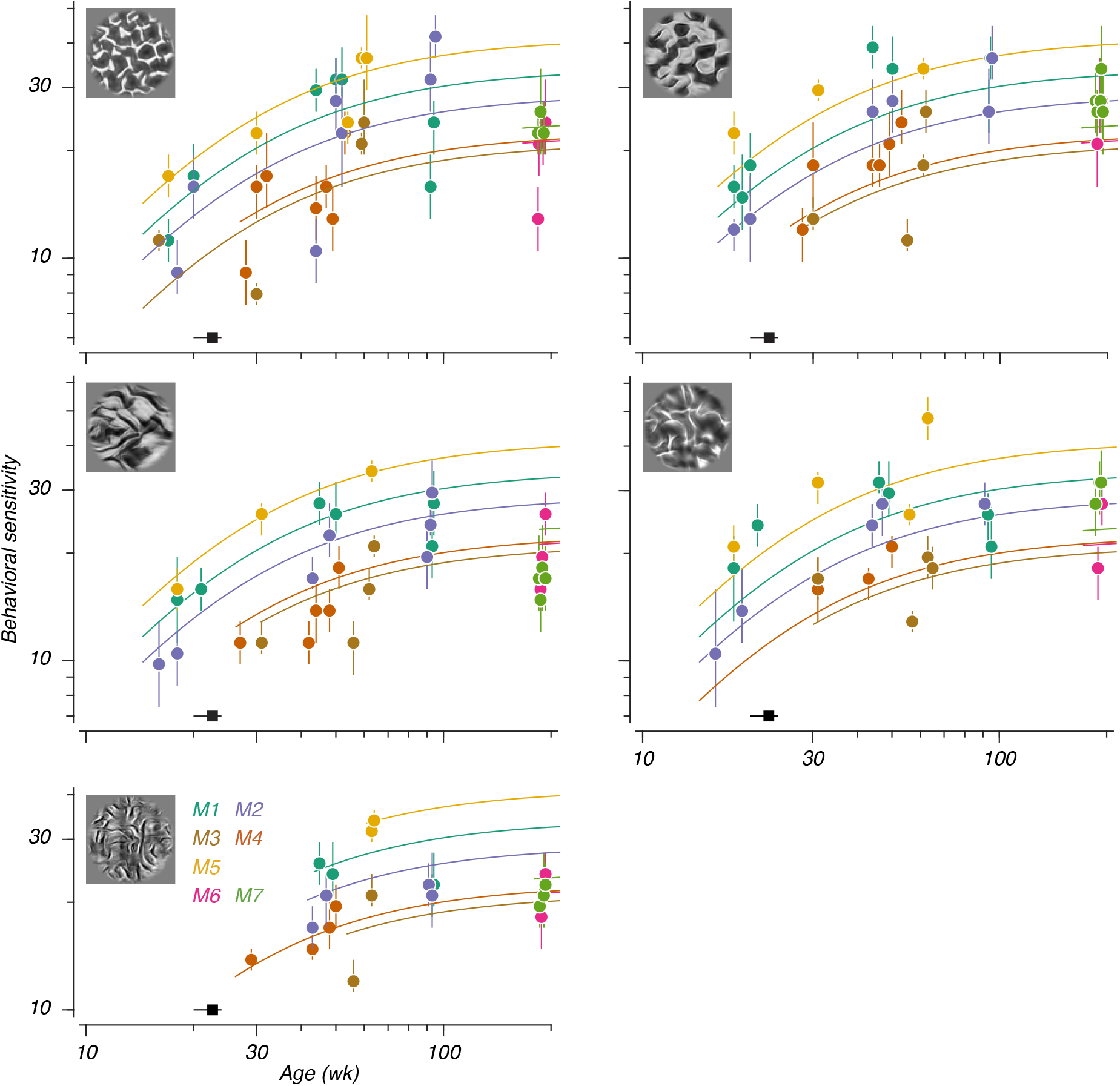
Behavioral texture sensitivities, plotted separately by texture family. Note the bottom panel, which starts from an older age than the others. Conventions the same as in Figure 2C.

**Supplementary Figure 2:**
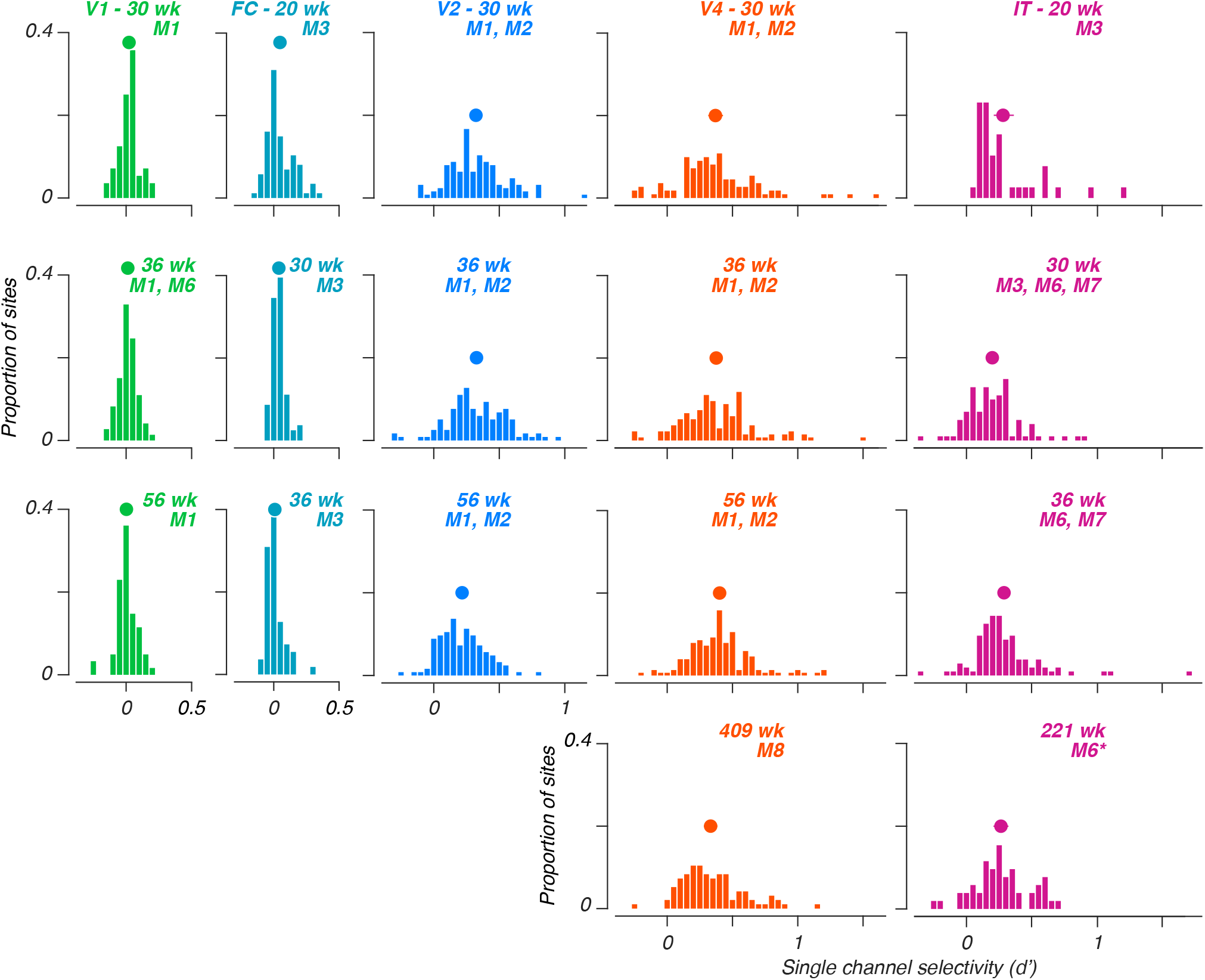
Single site texture selectivity distributions. Distributions of selectivity values, across areas. Each value reflects the mean selectivity for that site, across all texture families. Points represent distribution means, error bars represent 95% CIs around those means. Each subpanel represents data from a given age (note that ages are not always matched within a row). Monkeys used in each sample (see Table 1) are indicated with “M#” (we implanted M6 a second time in the opposite hemisphere, indicated with an asterisk).

**Supplementary Figure 3:**
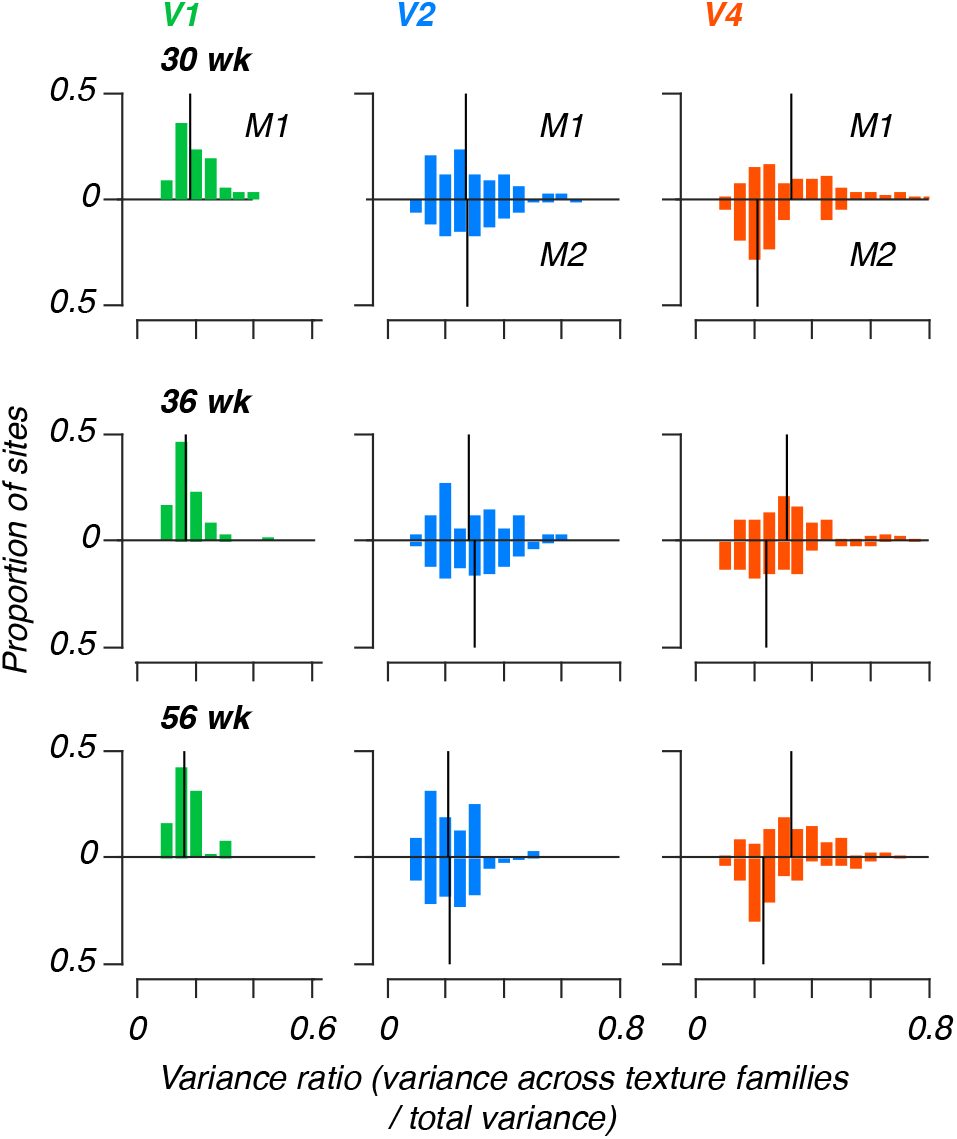
Variance ratios measured for longitudinal data. Each histogram depicts the distribution of variances across texture families (naturalistic and noise) to overall stimulus-driven variance, by age, by animal, for the two animals (M1 and M2) with the longest-lasting arrays.

**Supplementary Figure 4:**
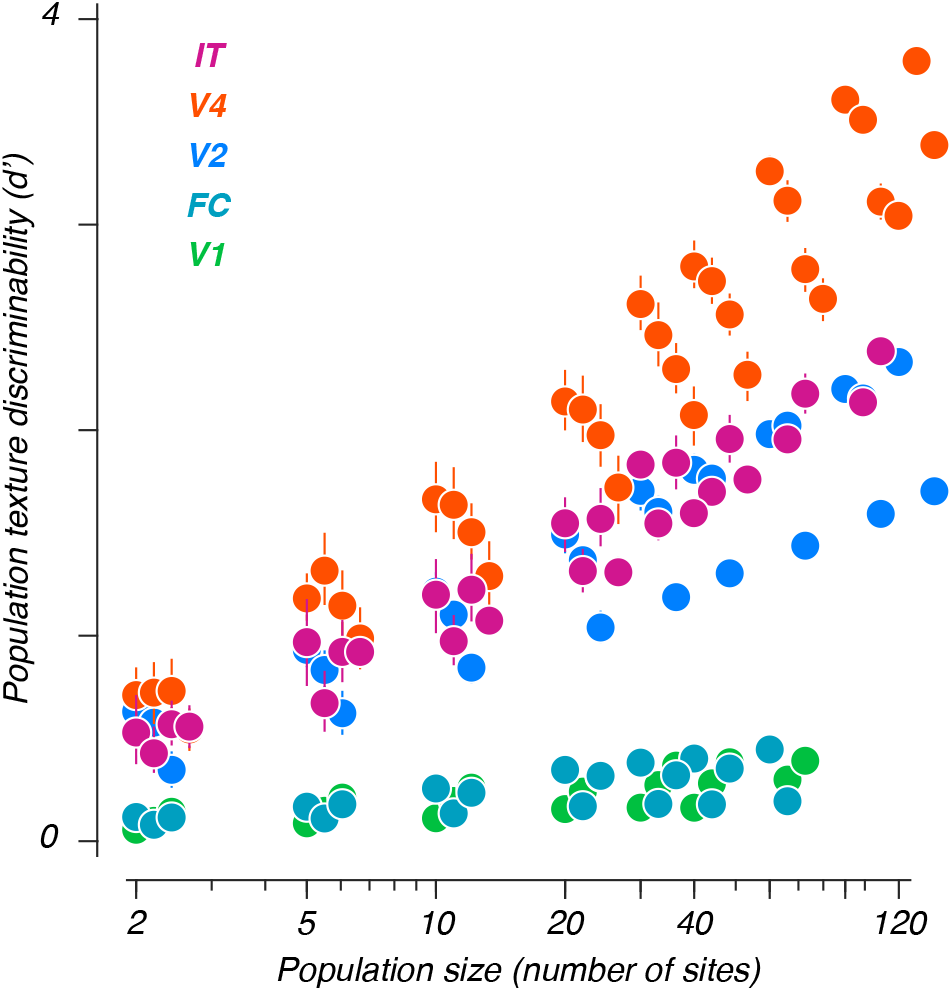
Performance scales with population size. Discriminability between naturalistic and noise textures versus population size. Each point represents data from a given area and age (for a given population size, data recorded at older ages are displaced increasingly to the right). Error bars depict 95% CIs across population resamplings; they are often smaller than the symbols, and thus invisible.

**Supplementary Figure 5:**
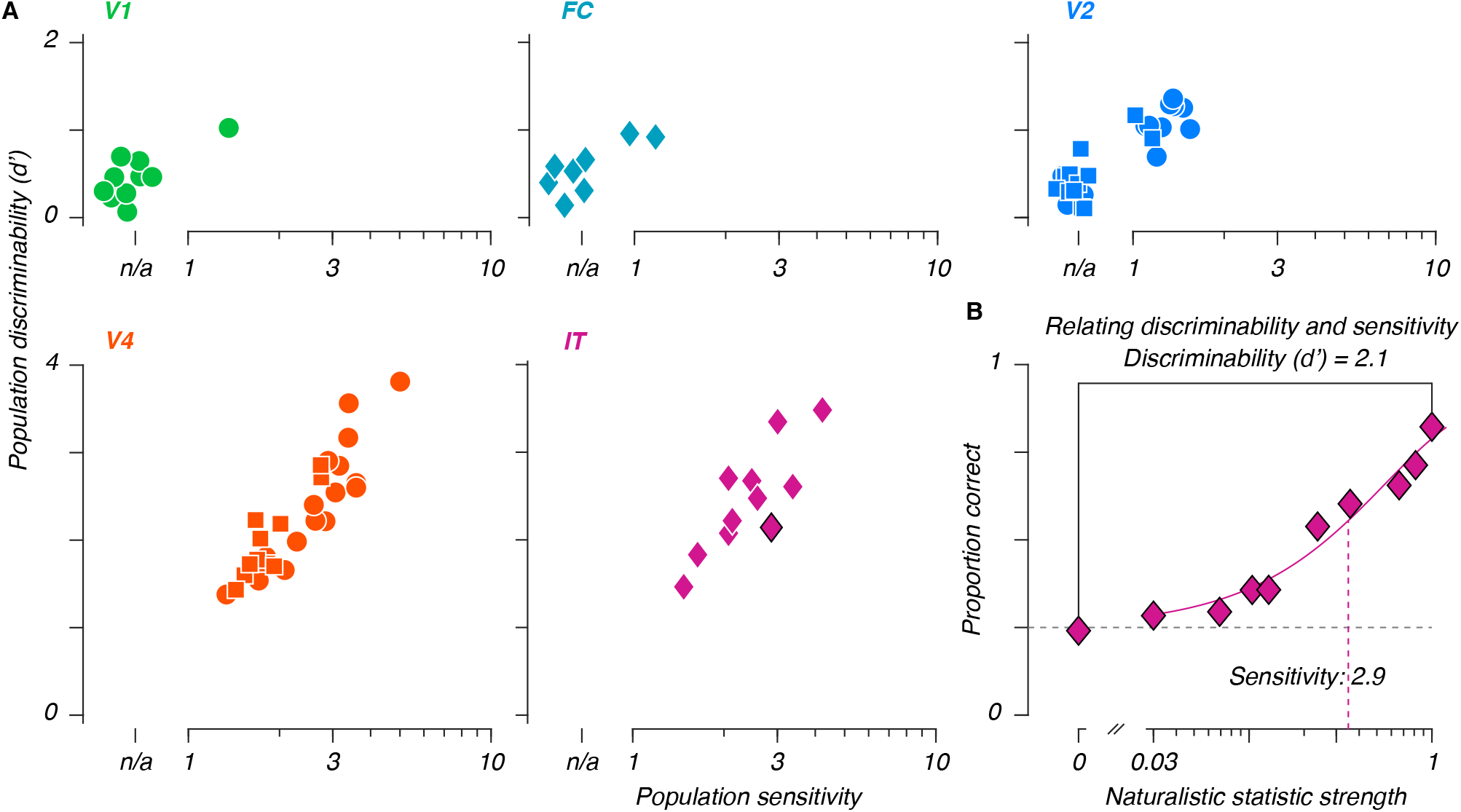
Comparing population discriminabilities and sensitivities. A: Discriminability between fully naturalistic and noise textures, versus neurometric sensitivities extracted from the same recordings. A discriminability can always be measured. Sessions for which a sensitivity could not be measured are offset to the left (n.s. – no sensitivity). B: Distinguishing sensitivity from discriminability – whereas sensitivity reflects performance across all levels of naturalistic structure, discriminability is limited to the distance between the noise and fully naturalistic texture endpoints (data reproduced from Figure 6A, this example corresponds to the point in panel A with a black rim).

